# Why diabetes matters in dementia studies: Excluding diabetes status masks regional mitochondrial DNA copy number changes in human hippocampus, amygdala, and cerebellum in Alzheimer’s disease

**DOI:** 10.64898/2026.06.03.729204

**Authors:** Aakruti Kaikini, Anqi Shi, Paul Francis, Russell Swerdlow, Claire Troakes, Jaimee Kennedy, Alan Hodgkinson, Afshan N. Malik

**Affiliations:** King’s College London, Diabetes and Obesity, School of Cardiovascular Medicine and Metabolic Sciences, Faculty of Life Sciences and Medicine.; University of Exeter, Dept of Clinical and Biomedical Sciences, University of Exeter Medical School, United Kingdom; University of Kansas Alzheimer’s Disease Research Center, University of Kansas School of Medicine, USA; King’s College London, London Neurodegenerative Diseases Brain Bank, Department of Clinical and Basic Neuroscience, Institute of Psychiatry, Psychology &Neuroscience; King’s College London, Department of Medical and Molecular Genetics, School of Basic and Medical Biosciences.

**Keywords:** Mitochondrial DNA, Diabetes, Alzheimer’s disease, post-mortem brain, co-morbidity, biobank

## Abstract

**INTRODUCTION:** Diabetes is a major risk factor for Alzheimer’s disease (AD), and both diseases involve mitochondrial dysfunction. We hypothesised that AD is associated with reduced mitochondrial DNA copy number (mtDNA-CN) in vulnerable brain regions, and that diabetes modifies these changes.

**METHODS:** Post-mortem hippocampus, amygdala, and cerebellum samples (N=66-77) from non-cognitively impaired (NCI) and AD donors, with and without diabetes, were analysed. mtDNA-CN was quantified by absolute quantification.

**RESULTS:** Overall, mtDNA-CN was lower in AD. However, stratification by diabetes revealed opposite changes: non-diabetic AD cases showed reduced mtDNA-CN, whereas diabetic cases showed higher mtDNA-CN across all regions irrespective of cognitive status.

**DISCUSSION:** These findings confirm multiregional loss of mtDNA-CN in the AD brain, most evident in the absence of diabetes. The functional significance of higher mtDNA-CN in the diabetic brain remains unclear, but evidence that diabetes can mask effects has important implications for dementia studies.

## 1. BACKGROUND

Alzheimer’s disease (AD), the leading cause of dementia, affects >57 million people worldwide, is predicted to rise to >150 million by 2050^1^. AD is increasingly recognised as a heterogeneous disorder, with multiple contributing mechanisms and more than 14 known modifiable risk factors^2,3^. Diabetes mellitus, a metabolic disorder characterised by chronic hyperglycemia, affects >550 million individuals worldwide, increases the risk of developing AD by 40-60%^4,5^.

Diabetes and AD share several pathological features including insulin resistance, mitochondrial dysfunction, chronic inflammation, and oxidative stress, suggesting that diabetes may accelerate neurodegenerative changes^6,7^. In addition to shared mechanisms, structural and molecular differences have been reported between diabetic and non-diabetic AD^8–10^. PET studies show a distinct clinical profile in dementia patients with diabetes, characterised by reduced amyloid burden and prominent tau pathology^11^. This suggests that clinically defined AD cohorts may comprise metabolically distinct subgroups with differing vascular, inflammatory, and bioenergetic contributions, potentially requiring tailored therapeutic strategies.

The human brain has high energy needs, largely met through oxidative phosphorylation via the electron transport chain, whose critical subunits are encoded by mitochondrial DNA (mtDNA). MtDNA is widely used as a biomarker of mitochondrial content, we have previously shown that some human brain regions can have >5000 mtDNA copies per cell (mtDNA-CN), making it a highly abundant molecule^12^. Changes in mtDNA-CN can influence energy metabolism, cellular signalling, and epigenetic programming, thereby affecting cellular health^13^. Reduced mtDNA-CN has been reported in several metabolic diseases, largely in the periphery, including diabetes, AD and cardiovascular disease^14–17^. Lower mtDNA-CN is generally associated with higher disease risk, supporting the idea that mtDNA-CN depletion may result in compromised mitochondrial function, loss of bioenergetic flexibility and cellular damage^18,19^.

Studies examining mtDNA-CN in the post-mortem human AD brain have produced inconsistent results. Some report reduced mtDNA-CN in cortical and hippocampal regions, with cerebellum typically unchanged. Whereas, large scale studies found no changes, leaving unresolved whether AD brain shows regional or general loss of mitochondrial content^20–25^. Importantly, most studies did not report the diabetic status of the donors, often due to incomplete clinical records. A previous study from our group is the only published study examining diabetic and non-diabetic AD^12^. We observed reduced mtDNA-CN in the parietal cortex of non-diabetic AD, with no change in frontal cortex or cerebellum. Unexpectedly, diabetic donors exhibited higher mtDNA-CN regardless of cognitive status in all 3 regions^12^. This contrasted with our original expectation that diabetes would exacerbate AD-related mitochondrial changes and instead suggested that diabetes and AD may exert opposing effects on brain mtDNA-CN.

Given the high prevalence of diabetes, affecting 28.8% of adults aged ≥65 years and reported in 6.0%–34.6% among individuals with AD, diabetes is likely a common comorbidity in AD^26–28^. Therefore, confirming whether regional or global loss of brain mitochondrial content is implicated in AD, and whether diabetic AD shows the same or different trends is important, as any differences may have implications for treatment strategies.

AD affects different brain regions, and vulnerability and clinical manifestations vary by region. For this study, we selected three brain regions with differential relevance to AD and diabetes: the hippocampus^29^, one of the earliest regions impacted and essential for spatial memory and learning; the amygdala^30^, which undergoes early atrophy linked to emotional and neuropsychiatric symptoms; and the cerebellum^31,32^, traditionally considered spared in AD but affected by diabetes and increasingly recognised for roles in cognition and insulin signalling. Whilst prior studies report mtDNA-CN reductions in human hippocampus and no changes in cerebellum, mtDNA-CN has never been measured in human amygdala. Furthermore, aside from diabetic cerebellar mtDNA-CN reported in our earlier study^12^, mtDNA-CN in hippocampus and amygdala of diabetic individuals has never been reported.

We therefore hypothesised that AD is associated with region-specific loss of mitochondrial content, measured as mtDNA-CN, primarily affecting the early-vulnerable hippocampus and amygdala. By stratifying donors according to AD and diabetes status, we aimed to determine whether diabetes modifies mtDNA-CN in a way that may help explain heterogeneity within AD.

## 2. METHODS

### 2.1. The aims, design, and setting of the study

This cross-sectional lab-based study aimed to quantify mtDNA-CN in three post-mortem human brain regions: hippocampus, amygdala and cerebellum, and determine if diabetes and AD, separately or in combination, is associated with altered mtDNA levels.

### 2.2. Ethical approval and informed consent

Project and ethical approval were obtained from the London Neurodegenerative Diseases Brain Bank (REC Reference 23/WA/0124). Post-mortem tissue was handled, stored, used, and disposed as per the Human Tissue Act 2004. Informed consent for the use of these samples for research was obtained by the brain bank.

### 2.3. Human post-mortem brain specimens

Samples for this study were selected based on the following predefined inclusion and exclusion criteria.

#### Inclusion criteria

- Confirmed diagnosis or absence of AD based on combined clinical and neuropathological assessment.
- Confirmed diagnosis or absence of diabetes mellitus based on documented clinical assessment.
- Availability of frozen tissue suitable for molecular analysis from the same donor.

#### Exclusion criteria

Samples were excluded if they showed significant history of other types of dementia including Parkinson’s disease, frontal temporal dementia or vascular dementia. All final group allocations and inclusion/exclusion decisions were made prior to laboratory analysis and mtDNA-CN was determined blinded to prevent bias.

### 2.4. Brain region sampling and processing

Samples had been processed according to the London Neurodegenerative Diseases Brain Bank brain bank’s standard anatomical dissection procedures, applied uniformly across donors. Hippocampal tissue was specifically from the CA1 subregion, whereas subregion information for amygdala and cerebellum was not recorded. Hippocampal and amygdala tissue was obtained from 77 donors, while cerebellar tissue was available for 66 donors. All tissues were fresh-frozen shortly after dissection and stored at −80°C until DNA extraction, with freeze–thaw cycles minimized to preserve integrity. Table 1 summarises demographic information of cases used in the study.

**Table 1:** Summary of Demographic Information for Included Cases.

| **Diabetes status** | **No** | **Yes** | **Total cohort** |
| --- | --- | --- | --- |
| Subjects (n) | 40 | 37 | 77 |
| Age (years)<sup>a</sup> | 79.0 ± 7.97 | 80.2 ± 11.15 | 79.1 ± 9.57 |
| PM delay (hours)<sup>a</sup> | 46.6 ± 20.4 | 44.6 ± 22.1 | 45.7 ± 21.0 |
| Sex (Female: Male) | 22:18 | 19:18 | 41:36 |
| Diagnosis (NCI:AD) | 20:20 | 18:19 | 38:39 |

<sup>a</sup>Values shown are mean ± standard deviation.

### 2.5 Quantification of mitochondrial DNA copy number

Total DNA was isolated using the QIAamp Fast DNA Tissue Kit, (Qiagen, UK), using the manufacturer’s protocol optimised for brain tissue as follows. Tissue (approximately 20 mg) was homogenised in the buffer using a Tissue Lyser II (Qiagen, UK) for 30 sec at 24 MHz. Homogenates were incubated in a thermomixer (Eppendorf™, UK) at 1000 rpm for 10 min at 56 °C with proteinase K. The resulting lysate was applied to the column and subjected to a series of washes with the provided wash buffers. DNA was eluted (50 µl elution buffer), concentration was determined by NanoDrop (Labtech International, UK), adjusted to a concentration of 10 ng/µl, and used in triplicate as a template with primer sequences targeting a unique region of human mtDNA (hMito) and the human nuclear gene beta-2-microglobulin (hB2M) (see Table S1 for primer sequences). Each 10 µl reaction consisted of 8 µl Master Mix (5 µl 2x Quantifast SYBR Green Master Mix, Qiagen; 500 nM forward and reverse primer; made up to volume with RNAase-free water) and 2 µl DNA template. Samples were loaded onto a 96-well plate in triplicate alongside a standard curve consisting of a serial dilution of 10^7^–10^3^ copies/2 µl of primer-specific PCR amplicons. All reactions were performed using the LightCycler 96 Real-Time PCR System (Roche Diagnostics, Switzerland). Absolute mtDNA-CN was calculated using the standard curve as a ratio of mitochondrial (hMito) to nuclear (hB2M) targets, representing cellular mtDNA content as described previously (MtDNA-CN), ^12,33^.

### 2.6. Statistical analysis

Statistical analyses were performed using GraphPad Prism software (Version 10.3.0) and R (Version 4.4.3).

For each brain region, outliers were defined using the interquartile range (IQR) method. The first quartile (Q1) represents the 25^th^ percentile of the data, and the third quartile (Q3) represents the 75^th^ percentile. The IQR is calculated as Q3 – Q1. Values greater than Q3 + 3×IQR or less than Q1 – 3×IQR were classified as extreme outliers.

Group comparisons were performed using one-tailed Student’s t-test when equal variances were confirmed by F-test or Welch’s t-tests when variances were unequal. Significance thresholds used after correcting for multiple testing were: *p< 0.0167; ** p< 0.0033, P < 0.00033. Sex differences in mtDNA-CN were assessed using an independent *t*-test, and Pearson correlation analyses were used to evaluate associations of mtDNA-CN with age and post-mortem interval (PMI).

We also performed inter-regional analysis in the 64 donors with complete dataset available for hippocampus, amygdala and cerebellum. Relationships between pairs of brain regions were examined using Spearman correlation analysis. Partial Spearman correlations were also performed to determine whether these pairwise relationships remained after controlling for the third brain region. To characterise multiregional patterns of mtDNA-CN, hierarchical clustering was applied to z-scored mtDNA-CN values using Ward’s minimum variance method. The resulting data-driven clusters were designated as low, moderate, or very high, and samples were arranged according to cluster membership for visualisation. Differences in cluster distribution across the four disease groups were assessed using a chi-square test of independence. Principal component analysis (PCA) was performed on z-score standardised mtDNA-CN values to identify the main patterns of variation across the three regions. Intraclass correlation coefficient (ICC) analysis was used to assess the consistency of mtDNA-CN across the three regions within the same individual. Kendall’s coefficient of concordance (W) was used to determine whether the relative order of the three brain regions was similar across individuals. These analyses were first carried out in the whole dataset and then repeated after stratification into the four disease groups.

## 3. RESULTS

The study utilised post-mortem brain samples from 3 regions of the human brain, namely hippocampus, amygdala, and cerebellum. Hippocampus and amygdala were available for all 77 cases, whereas cerebellum was available for 66 cases. Subjects were categorized as having no cognitive impairment (NCI, n = 38) or AD (n = 39), based on in-life diagnosis and/or Braak neuropathology^34^. Using their cognitive and diabetes status, the subjects were further categorized into 4 groups; No diabetes, non-cognitively impaired with a Braak score of 0-2 (ND-NCI, n=20); Diabetes and non-cognitively impaired with a Braak score of 0-2 (D-NCI, n=18); No diabetes, Alzheimer’s disease with a Braak score of 5-6 (ND-AD, n=20); Diabetes and Alzheimer’s disease with a Braak score of 5-6 (D-AD, n=19) (Table 1).

As determined by the IQR method described above, two mtDNA-CN values for cerebellum (452.0 and 704.6) exceeded the defined outlier thresholds (Q1 – 3×IQR or Q3 + 3×IQR) and were therefore excluded from subsequent analyses resulting in n=64 for cerebellum.

### 3.1. Regional variation in mtDNA-CN in the 3 brain regions

We observed regional differences in mtDNA-CN across the three brain regions examined (Figure 1A). The amygdala showed the highest mtDNA-CN, ranging from ∼ 200 to 1,100 copies per cell (mean 503 copies/cell), followed by the hippocampus, which ranged from about 63 to 1,110 copies per cell (mean 421 copies/cell). The cerebellum had the lowest mtDNA-CN, ranging from ∼38 to 376 copies per cell (mean 160 copies/cell). In the whole dataset (n=64), the dominant regional hierarchy was Amygdala>Hippocampus>Cerebellum, observed in 40 out of 64 individuals (62.5%), followed by Hippocampus> Amygdala> Cerebellum in 21 out of 64 individuals (32.8%) (Figure 1B). Spearman correlation analysis of the whole dataset showed a moderate positive correlation between hippocampus and amygdala (ρ = 0.48, p = 0.0001) and a weaker but significant positive correlation between hippocampus and cerebellum (ρ = 0.35, p = 0.005). In contrast, the correlation between amygdala and cerebellum was weak and did not reach significance (ρ = 0.24, p = 0.053) (Figure 1C). To further examine whether these relationships were direct or indirect, partial correlation analysis was performed (Figure 1C). The hippocampus-amygdala correlation remained strong after controlling for cerebellum (partial ρ = 0.44, p = 0.0003), and the hippocampus-cerebellum correlation also remained significant after controlling for amygdala (partial ρ = 0.28, p = 0.029). However, the amygdala-cerebellum correlation was lost after controlling for hippocampus (partial ρ = 0.09, p = 0.48). These findings suggest that hippocampus has independent relationships with both amygdala and cerebellum, whereas the apparent amygdala-cerebellum association is largely explained by their shared relationship with hippocampus.

**Figure 1.**
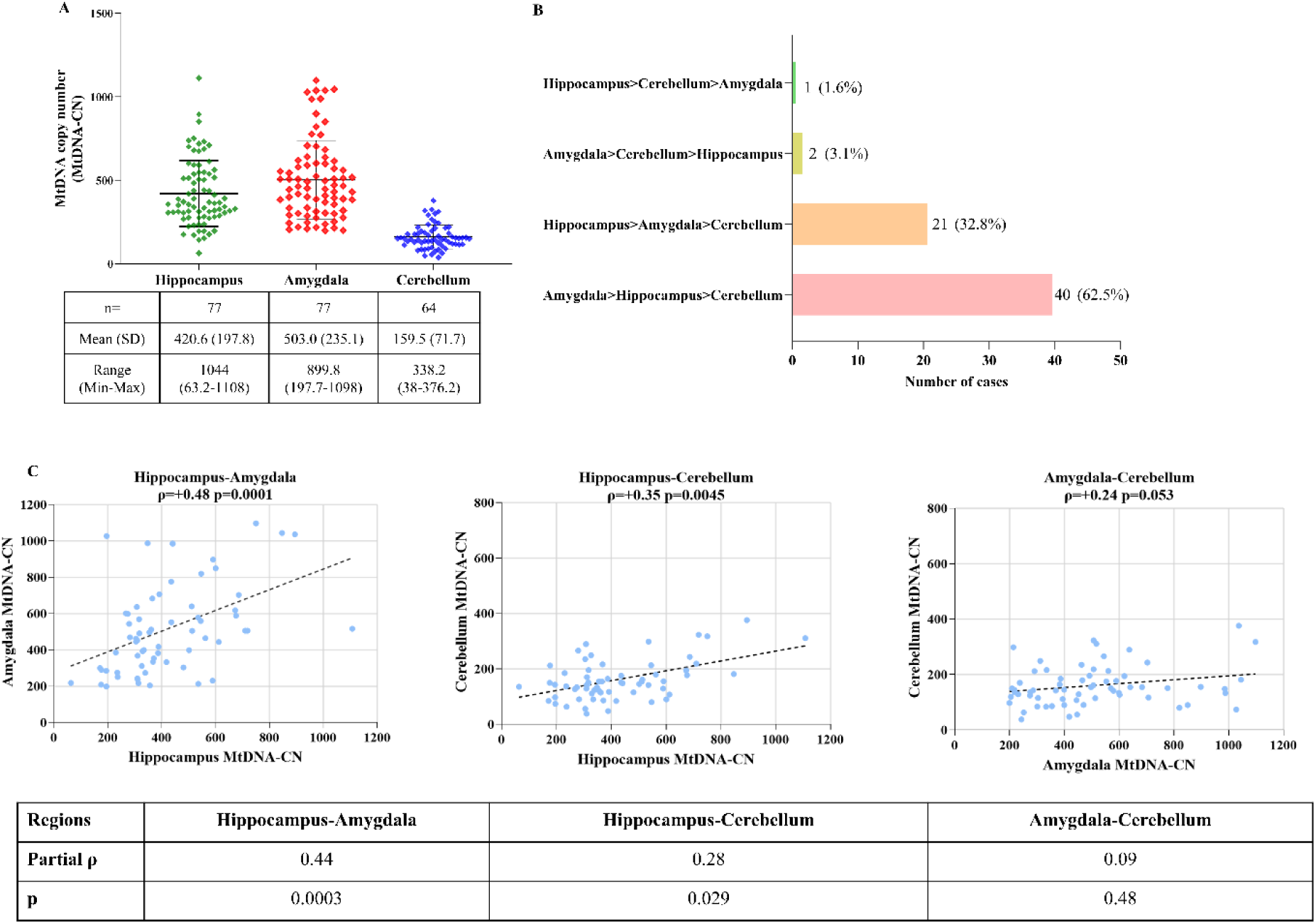
Regional differences and correlation between mtDNA-CN in three regions of human post-mortem brain. **(A)** MtDNA copy number (mtDNA-CN) was measured in hippocampus (n=77), amygdala (n=77) and cerebellum (n=64) of the human post-mortem brain using absolute quantification by real-time qPCR, with values expressed as ratio of mitochondrial to nuclear genome. Each datapoint represents mtDNA-CN for an individual case, with data shown as mean ± SD. Group sizes (n) and descriptive data (mean, standard deviation, range, minimum and maximum) shown in the table below the graph. **(B)** Patterns of regional mtDNA copy number levels in hippocampus, amygdala, and cerebellum within the same individual. The panel displays the frequency of the four possible ranking orders, indicating which region has higher or lower mtDNA-CN relative to the others. **(C)** Correlation of mtDNA-CN between brain region pairs. Scatterplots show pairwise comparisons of mtDNA-CN between hippocampus-amygdala, hippocampus-cerebellum, and amygdala-cerebellum. Spearman’s correlation coefficient (ρ) and corresponding p value are shown on each graph. A linear regression line is included for visualisation. Partial correlation analysis, controlling for mtDNA-CN in the third region, was also performed for each region pair, with partial ρ and p values shown in the table below. Correlations involving hippocampus remained significant after adjustment, whereas the amygdala-cerebellum association was not significant.

### 3.2. mtDNA-CN shows no correlation with age, post-mortem interval (PMI), and sex

MtDNA-CN was compared between males and females in each brain region using an independent Student’s *t*-test. No significant sex differences were observed in any of the regions examined (Figure 2A). Pearson correlation analyses were then conducted to examine the relationship of mtDNA-CN with age and PMI in the amygdala, hippocampus, and cerebellum. No significant correlations were detected between mtDNA-CN and either age (Figure 2B) or PMI (figure 2C) in any brain region. Together, these findings indicate that sex, age, and PMI did not influence mtDNA-CN levels in this study.

**Figure 2.**
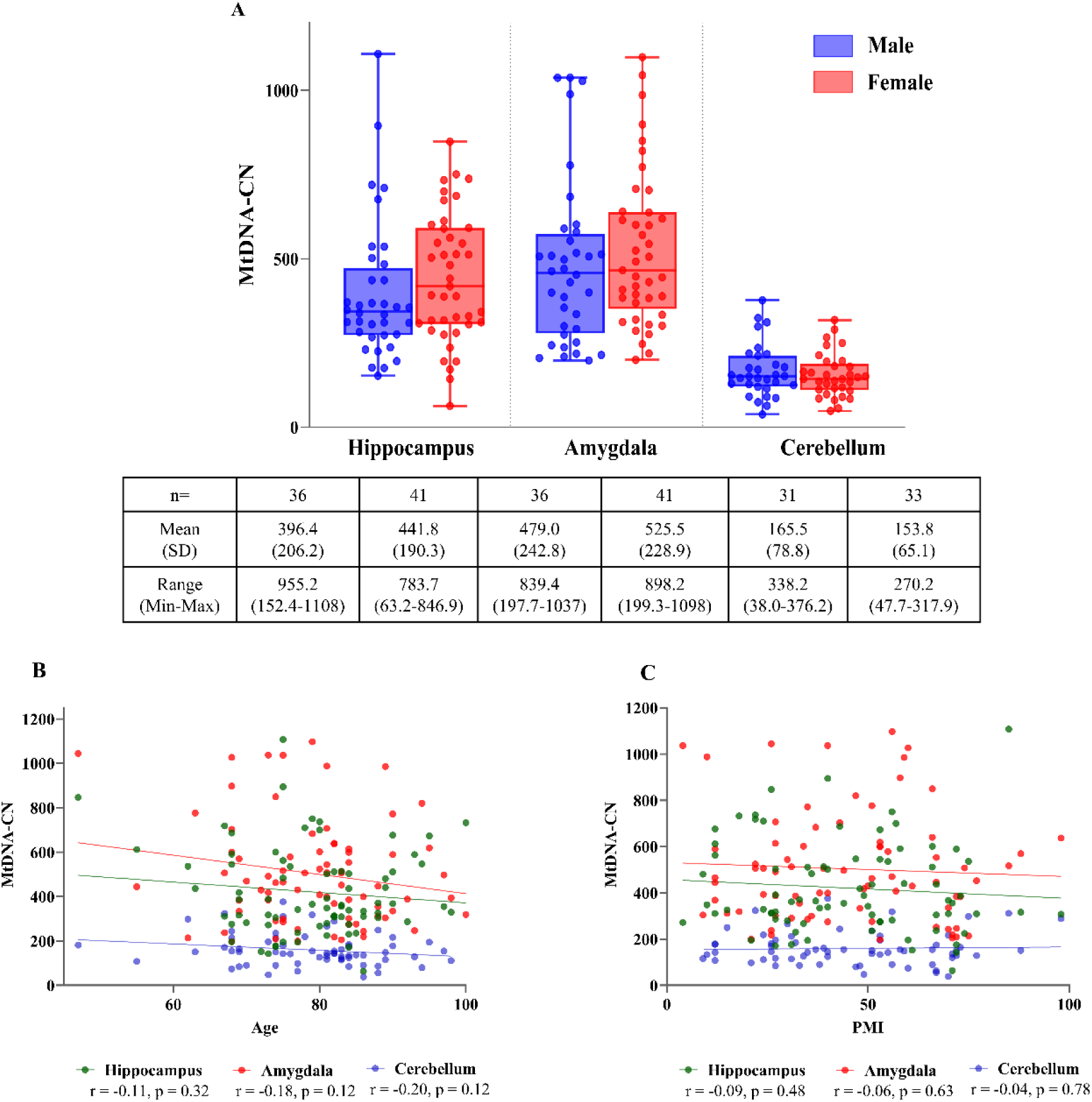
Sex, age, and post-mortem interval do not influence mtDNA copy number in human brain. mtDNA-CN in hippocampus (n=77), amygdala (n=77), and cerebellum (n=64) was examined in relation to (A) biological sex, (B) age, and (C) post-mortem interval in each of the three brain regions. **(A)** mtDNA-CN across the three brain regions is stratified by sex (blue, male; red, female). Each dot represents mtDNA-CN for an individual sample; boxplots display the median and interquartile range. No significant differences were observed between males and females in any brain region (independent t-test). **(B–C)** Associations between mtDNA-CN and either age **(B)** or post-mortem interval (PMI) **(C)** across all samples are shown. Each dot represents mtDNA-CN for an individual sample, with colours indicating brain region as shown. Pearson correlation coefficients (r) and corresponding p values are shown on each panel, and a simple linear regression line is included for visualisation No significant associations were observed between mtDNA-CN and either age or PMI in any brain region (p > 0.05).

### 3.3. mtDNA-CN is reduced in AD and increased in diabetes

In all three brain regions, comparison of AD subjects with NCI showed modest reductions of 16-19% of mtDNA-CN, reaching significance in nominal P values for two brain regions (hippocampus and amygdala, P<0.05, Fig. 3A, Table 2) which did not survive after correcting for the number of brain regions tested (P>0.0167 in all regions). The mean AD to NCI ratio across all three regions was ∼0.83, indicating a similar magnitude of reduction in each region and suggesting a biologically consistent effect.

**Figure 3.**
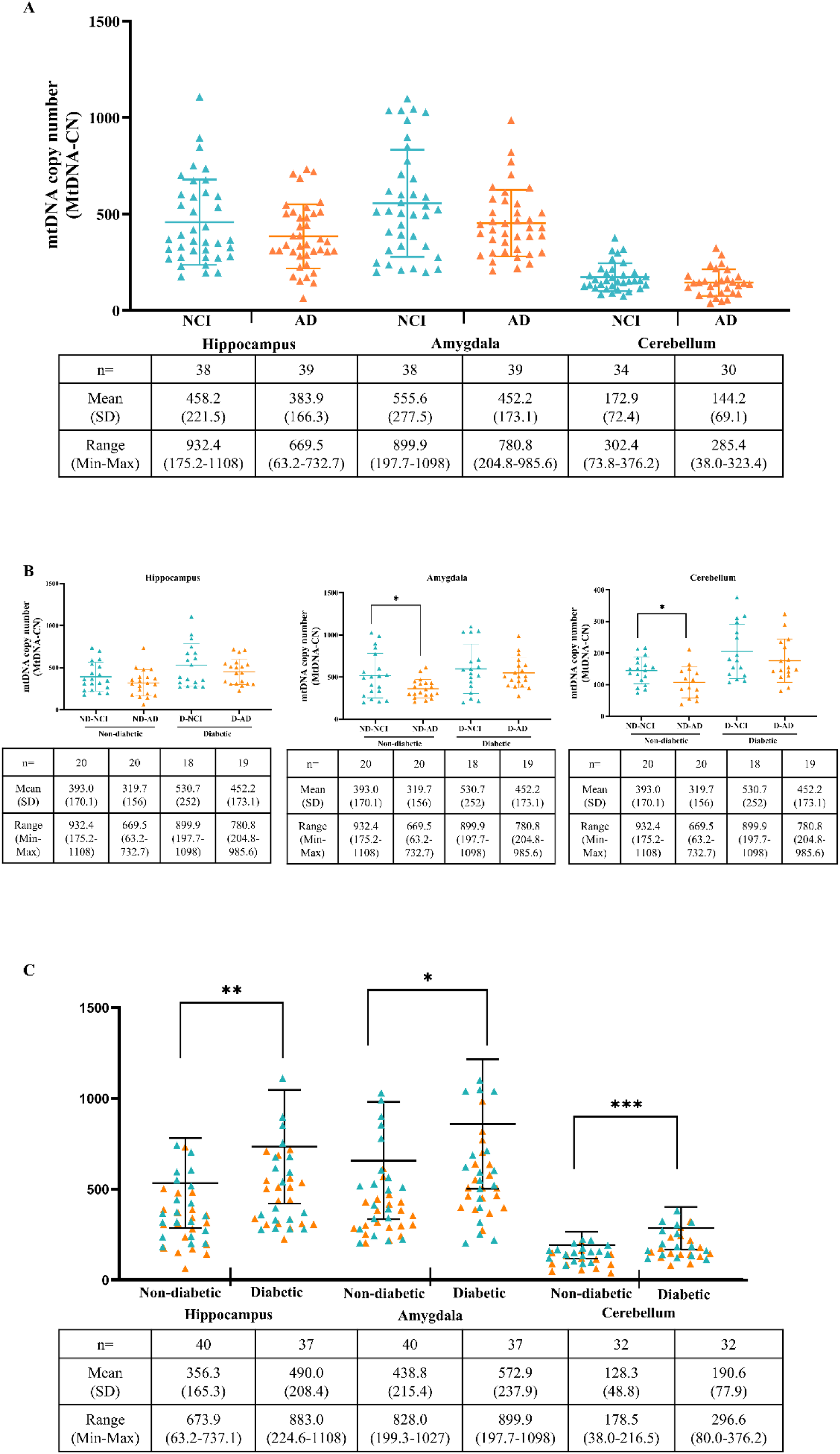
Mitochondrial DNA copy number in three regions of human brain changes in Alzheimer’s disease and diabetes. mtDNA-CN in **(A)** AD vs NCI **(B)** NCI and AD with (D-NCI, D-AD) and without (ND-NCI, ND-AD) diabetes **(C)** non-diabetes vs diabetes. NCI shown as green triangles, AD shown as orange triangles. Each point represents the mtDNA-CN of an individual sample. Group sizes (n) and descriptive data (mean, standard deviation, range, minimum and maximum) shown as tables below each figure panel. P-values were determined using student t-test when variances were equal or Welch’s t-test when variances were unequal. To control for multiple testing across three brain regions, the nominal P value threshold (<0.05) was divided by 3. Significance thresholds after correcting for multiple testing: *p< 0.0167; ** p< 0.0033, P < 0.00033.

**Table 2.** Summary of P values for each comparison for the 3 brain regions investigated.

| **Comparison** | **P-values** |  |  |
| --- | --- | --- | --- |
|  | **Hippocampus** | **Amygdala** | **Cerebellum** |
| NCI vs AD (Figure 3A) | 0.0498 | 0.0281<sup>#</sup> | 0.0555 |
| Non-diabetic NCI vs non-diabetic AD (Figure 3B) | 0.0819 | 0.0113<sup>#</sup> | 0.0159 |
| Diabetic NCI vs diabetic AD (Figure 3B) | 0.1307<sup>#</sup> | 0.2266<sup>#</sup> | 0.1526 |
| Non-diabetic vs diabetic (Figure 3C) | 0.0012 | 0.0057 | 0.0002<sup>#</sup> |

Group comparisons were carried out using either student t-test (when variances were equal)
or Welch's t-test (when variances were unequal, values marked with #).

However, when restricting analysis to non-diabetic subjects, stronger reductions of between 19-30% were observed in ND-AD cases compared to ND-NCI cases (Fig 3B, Table 2) with mean ratios of ∼0.81 in hippocampus, 0.70 in amygdala and 0.74 in cerebellum. Two brain regions, amygdala and cerebellum, reached statistical significance even after correction for multiple testing despite smaller sample sizes (P<0.0167, Table 2). By contrast, in subjects with diabetes, D-AD cases showed only small reductions of 8-15% compared to D-NCI (in all regions, P>0.05, Fig 3B, Table 2).

Finally, and across the entire cohort, diabetes itself was associated with the opposing direction of change, showing highly significant increases in mtDNA-CN of 30-49% across the three regions which remained highly significant after correction for multiple testing (P<0.0167 in all regions, Fig 3C, Table 2). In individuals with diabetes, mean mtDNA-CN was 37.5% higher in hippocampus, 30.6 % higher in amygdala and 48.5% higher in cerebellum compared to the non-diabetic cases, corresponding to mean ratios of ∼1.38 in hippocampus (P = 0.0012), 1.27 in amygdala (P = 0.0057), and 1.49 in cerebellum (P < 0.0002), respectively.

Notably, AD alone was linked to lower mtDNA-CN (Fig.3B) whilst diabetes was linked to higher mtDNA-CN (Fig 3C), showing opposing effects that effectively cancelled each other out when diabetes status was not considered (Fig. 3A).

### 3.4 mtDNA-CN levels in different brain regions of the same individual

As we had used three brain regions from the same individuals (n=64, Figure 4A), we examined the similarities in mtDNA-CN levels across the 3 regions to determine any group patterns using hierarchical cluster analysis using ward’s method on z-scored mtDNA-CN. This resulted in identification of three distinct clusters: low, moderate and very high (Figure 4B). Cluster membership was found to be significantly associated with disease group (χ² = 24.2, df = 6, p = 0.0005): <u>The lower cluster</u> (n=23) was characterised by uniformly below average mtDNA-CN across all three regions (mean z-scores: hippocampus = −0.67, amygdala = −0.76, cerebellum = −0.69) and was enriched for ND-AD (11/23, 48%) and ND-NCI (7/23, 30%), with only 1 D-NCI (4%); <u>The moderate cluster</u> (n = 36) was characterised by near-average mtDNA-CN across all regions (mean z-scores: hippocampus = +0.16, amygdala = +0.39, cerebellum = +0.12) and was the largest cluster showing a balanced composition across all four disease groups (ND-NCI = 11, D-NCI = 11, D-AD = 11, ND-AD = 3); <u>The very high cluster</u> (n = 5) was characterised by markedly elevated mtDNA-CN, particularly in hippocampus and cerebellum (mean z-scores: hippocampus = +1.93, amygdala = +0.70, cerebellum = +2.31). This cluster was dominated by D-NCI s (4/5, 80%), with 1 D-AD and no non-diabetic donors.

**Figure 4.**
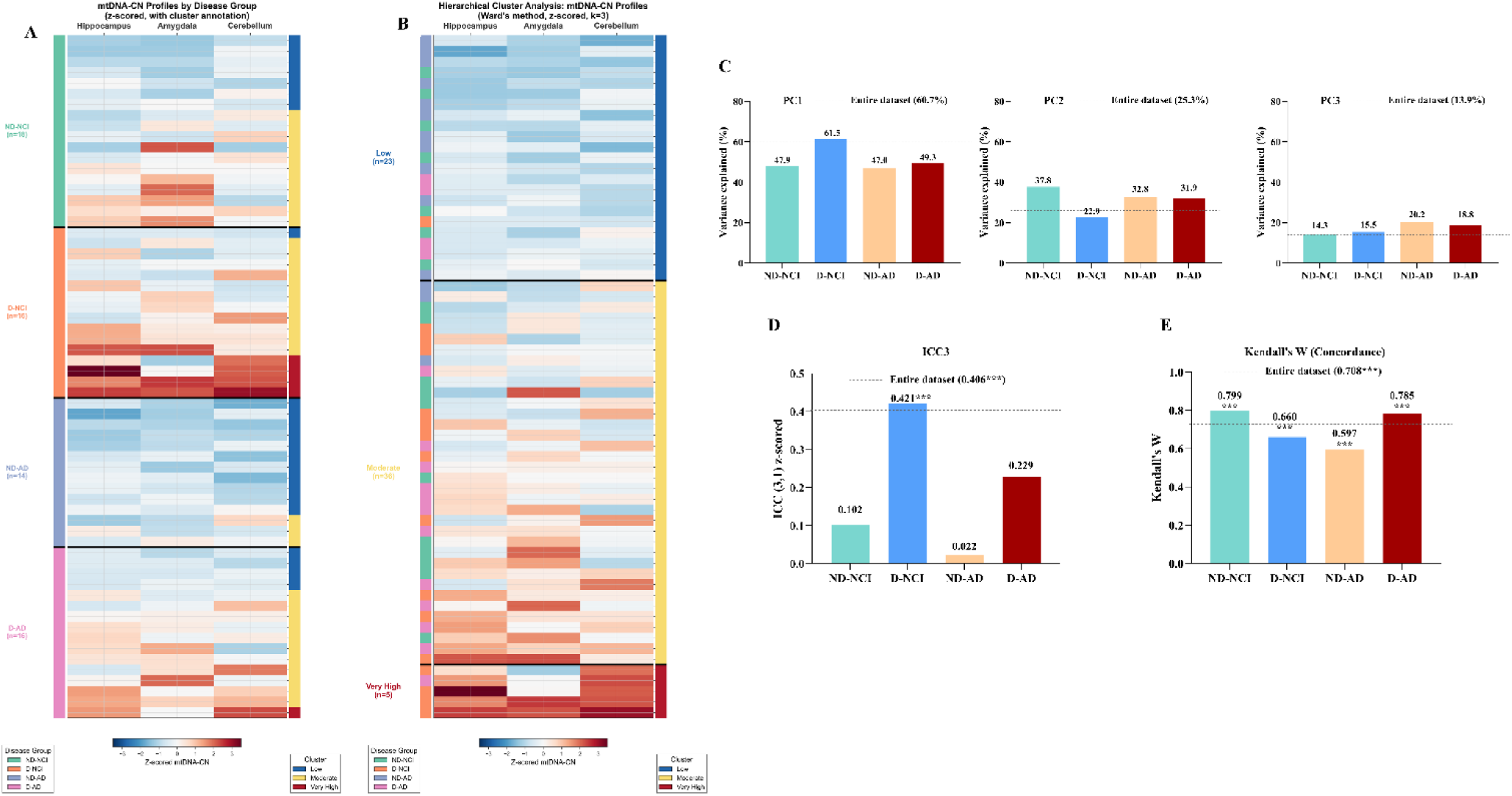
Regional distribution and intra-individual patterns of mitochondrial DNA copy number in hippocampus, amygdala and cerebellum. Levels of mtDNA-CN across the three brain regions in each individual were z-scored and are shown as heatmaps. **(A)** Donors are ordered by disease group (ND-NCI, D-NCI, ND-AD and D-AD). Within each group, donors are further ordered by hierarchical cluster assignment and mean z-score. The left colour bar indicates disease group, and the right colour bar indicates cluster assignment (blue = low, yellow = moderate, red = very high). Case IDs are shown on the right, and black lines separate the disease groups. **(B)** Donors are ordered by hierarchical clustering using Ward’s minimum variance method (k = 3), identifying three mtDNA-CN profiles: low (n = 23), moderate (n = 36) and very high (n = 5). The low cluster was enriched for ND-AD cases (48%), whereas the very high cluster was dominated by D-NCI cases (80%). A chi-square test of independence showed that the distribution of cases across the three mtDNA-CN clusters differed significantly among the four disease groups (χ² = 24.2, df = 6, p = 0.0005). **(C)** The main patterns in the mtDNA-CN dataset across the three brain regions were examined using principal component analysis (PCA). PC1 reflected the dominant shared mtDNA-CN pattern across hippocampus, amygdala and cerebellum, whereas PC2 and PC3 captured smaller region-specific variation. The distribution of PC1, PC2 and PC3 scores across the disease groups is shown. The x-axis shows the disease groups, and the y axis shows the percentage of total variance explained by each component. Dashed lines indicate the corresponding variance explained by each component in the whole dataset. PC1 explained 60.7% of the total variance in the whole dataset and differed significantly between groups (Kruskal-Wallis H = 26.13, p < 0.0001), whereas PC2 and PC3 showed no significant group differences. **(D)** Intraclass correlation coefficient (ICC) was used to assess within-individual concordance in mtDNA-CN across hippocampus, amygdala and cerebellum. The whole dataset showed moderate within-individual concordance, but this was not uniform across groups. D-NCI showed the strongest concordance and was the only group with a significant ICC (p = 0.017). **(E)** Kendall’s W was used to assess the consistency of regional ranking in mtDNA-CN across hippocampus, amygdala and cerebellum in the whole dataset and within each disease group. Kendall’s W was significant in the whole dataset (W = 0.57, *p* = 0.0003) and remained significant in all four disease groups (p < 0.001).

Principal component analysis (PCA) of the whole dataset (Figure 4C) showed that the first principal component (PC1) accounted for 60.7% of the total variance, with all three brain regions loading positively and with similar magnitude (Table S2). This indicates that a large part of the variation reflects a shared pattern seen consistently across amygdala, hippocampus and cerebellum, where all regions tend to change together. PC2 explained an additional 25.3% of the variance and primarily captures divergence between amygdala and cerebellum, whereas PC3 (13.9%) reflected variation that was more hippocampus specific (Figure 4C, Table S2). Taken together, these components demonstrate that the dataset contains both a strong common mtDNA-CN pattern across regions and meaningful region-specific differences.

We next assessed whether this whole dataset structure was retained when donors were stratified into the four groups (Figure 4C, Table S2B-C). PC1 scores differed significantly across the four disease groups (Kruskal–Wallis H = 26.13, p < 0.0001, Table S2B), whereas PC2 and PC3 did not (Table S2B). Because PC1 was the dominant component and the only principal component that differed significantly across disease groups (Table S2B), group-specific loading analysis was focused on PC1. In D-NCI, PC1 explained 61.5% of the variance and all three regions loaded equally (Table S2C), indicating a robust shared pattern within this group. In contrast, PC1 explained a smaller proportion of variance in the remaining groups (47.0–49.3%), suggesting more heterogeneous within-group structure and weaker uniformity across regions (Figure4C, Table S2C). In both ND-NCI and ND-AD, hippocampus and amygdala loaded together on PC1 whereas cerebellum loaded in the opposite direction (Table S2C). D-NCI donors had the highest mean PC1 scores (+1.02 ± 1.66, Table S2B), while ND-AD donors had the lowest (−1.22 ± 0.57, Table S2B). These opposing shifts are consistent with diabetes being associated with **elevated** and AD with **reduced** mtDNA-CN across all three regions

Intraclass correlation (ICC) analysis of the full dataset showed **moderate within-individual consistency** across the three brains regions (ICC= 0.406, p < 0.001, Figure 4D, Table S3A). When examined separately by disease group (Figure 4D, Table S3A), D-NCI showed the **strongest regional concordance** (ICC3 = 0.423, p = 0.003) and was the only group with statistically significant within-individual agreement. In contrast, ND-AD showed virtually no concordance (ICC3 = −0.018, p = 0.524), indicating that mtDNA-CN levels in one region did not correlate with other regions in this group. Both ND-NCI and D-AD showed only weak and non-significant concordance (Figure 4D, Table S3A). These results indicate that the moderate ICC observed in the combined dataset does not reflect a uniform pattern across all disease groups.

In contrast, Kendall’s W which assesses the **consistency of regional ranking** rather than absolute agreement, was significant in the full dataset (W = 0.57, p = 0.0003) and remained significant within each of the four groups (Figure 4E, Table S3B). This indicates that the relative ordering of regions was generally preserved across individuals even when absolute concordance was weak.

## 4. DISCUSSION

Diabetes is a well-established risk factor for AD, yet it remains unclear whether diabetes accelerates canonical AD pathological pathways or contributes through independent mechanisms^2,5^. Although diabetes and AD are distinct diseases, they share intersecting biological processes, including chronic inflammation, oxidative stress and impaired insulin signalling in both the periphery and the brain, and mitochondrial dysfunction, which may converge to influence neurodegeneration^6,7,29^. Growing evidence suggests that diabetes-associated dementia risk cannot be fully explained by classical AD pathology alone, suggesting that diabetes may impose additional or alternative biological changes in the brain. This has led to the emerging view that AD may comprise of metabolically distinct subtypes, shaped by vascular, inflammatory and bioenergetic factors which need to be characterised to guide more precise therapeutic strategies^2^. To this end, in the current study, we examined mtDNA-CN levels across three regions of the human brain, amygdala and hippocampus, which are early and vulnerable targets of AD pathology^29, 30^, and cerebellum, which is thought to be relatively resistant^31^, to determine whether AD is associated with loss of mitochondrial content and if diabetes modifies regional mtDNA patterns in AD.

The first and central key finding of this study is that mtDNA-CN shows opposing directional changes in AD and diabetes and that this difference can impact pooled analyses results and mask AD only changes (Figure 3, Table 2). When we analysed mtDNA-CN levels without accounting for diabetes in the full data set (n=64-77), mtDNA-CN appeared modestly reduced with small region wide decreases that were nominally significant but did not withstand correction for multiple comparisons. However, once we removed the diabetic cases, the differences between ND-NCI and ND-AD mtDNA-CN became larger reaching significance for amygdala, and cerebellum, with effect sizes of similar magnitude in all three regions. This pattern suggests a robust biological difference in AD that is partially obscured when diabetes cases are included in the analysis. In contrast, within the diabetes groups, AD cases (D-AD) only showed minimal and non-significantly lower mtDNA-CN indicating that diabetes attenuates or masks the mitochondrial deficits typically associated with AD.

Direct comparison of diabetic versus nondiabetic donors further clarified this opposing pattern of brain mtDNA-CN levels in AD and diabetes (Fig. 3). Diabetes was associated with significantly higher levels of mtDNA-CN across all 3 brain regions. This consistent increase explains why AD effect seems weaker when diabetic donors are included in the analysis, and highlights diabetes as a strong modifier of mtDNA-CN in the brain. These findings strengthen the view that AD is associated with lower mtDNA-CN, whereas diabetes is associated with a higher mtDNA-CN, with the two conditions exerting counteracting cross regional influences that can cancel each other out in aggregated datasets. This is consistent with our previous report.^12^ Using a smaller and different cohort, we measured mtDNA-CN in frontal and parietal cortex and cerebellum, and our data showed similar trends, diabetes cases had higher mtDNA-CN in all regions whereas ND-AD was associated with lower mtDNA-CN in parietal cortex only. The current study extends these earlier findings and shows that these changes are not region specific.

The second key finding in our study is that although mtDNA levels vary between brain regions, as shown in both our earlier^12^ and current study, across the different brain regions, similar trends of diabetes or AD associated mtDNA-CN changes are seen, as all regions tended to change together in the presence of diabetes. Only D-NCI donors exhibited significant within-individual concordance across regions, while ND-AD donors showed no cross-regional agreement at all. Nevertheless, Kendall’s W indicated that the relative ordering of regions was preserved across all groups, even when absolute values diverged. Generally, the highest levels were found in amygdala (197.7-1098 copies/cell), slightly lower in hippocampus (63.2-1108 copies/cell), and significantly lower in cerebellum (38.0-376.2 copies/cell).

Our results indicate that diabetes acts as an effect modifier of mtDNA-CN patterns in AD brain. An implication of this finding is that diabetes should not be treated as a background comorbidity when interpreting mitochondrial parameters in AD brain tissue. In practical terms, pooled analyses that do not account for diabetes status will tend to underestimate AD-associated reductions, and the observed magnitude will depend on the metabolic composition of the cohort. This provides a plausible explanation for inconsistent mtDNA-CN findings across studies in AD human brain where differences in AD are often modest and not consistently detected across studies (Table S4). It is of note that diabetes status is not systematically incorporated as a covariate in many large-scale analyses, leaving the possibility that metabolic heterogeneity contributes to inconsistency in mtDNA-CN estimates^23^. Additionally, in the current study we used a matched design to balance AD and control participants with and without diabetes, but in the general population, diabetes is more prevalent in AD^27^. This imbalance likely amplifies confounding, where diabetes-driven increases in mtDNA-CN disproportionately occur in AD, masking or distorting AD-specific signals.

Neither age nor sex were associated with mtDNA-CN in the three brain regions investigated. This aligns with large-scale sequencing studies suggesting that mtDNA-CN largely appears uninfluenced by age and sex in post-mortem brain tissue. For instance, Rice et al (2014) reported no correlation between mtDNA-CN and age in either AD or control cohorts.^21^ Similarly, analyses from Klein et al. (2021) found no effect of age or sex on mtDNA-CN after accounting for regional neuropathology across multiple brain regions, including the dorsolateral prefrontal cortex, posterior cingulate cortex, and cerebellum.^23^ Wei et al. (2017) further highlighted disease-specific patterns, reporting no age association in AD but an age-dependent increase in Creutzfeldt–Jakob disease.^24^

A question arising from these findings is whether the higher and a co-ordinated pattern of mtDNA-CN seen in diabetic brain has a functional impact. Whilst the potential impact of the increased mtDNA-CN remains unknown, the higher brain mtDNA-CN seen in diabetic cases may not be functional and may not be associated with improved mitochondrial function but instead is suggestive of an alternative mtDNA mediated pathological process which could affect cellular health in an independent manner to AD associated pathologies. That disease associated mtDNA-CN increase may not always be functional has been suggested by our previous findings in diabetic models: we have shown that in cell models high-glucose exposure increases mtDNA-CN, but this rise does not translate into higher transcription or translation of mtDNA-encoded mRNAs, indicating that the increase in mtDNA-CN represents a compensatory response by stressed mitochondria rather than an improvement in function^7^, in fact in these studies, increases in mtDNA-CN which was not functional preceded bioenergetic deficit and cellular damage^7,19^. We reported a similar sequence of events in a mouse model of NAFLD and showed that an early maladaptive increase in hepatic mtDNA preceded mitochondrial dysfunction and liver damage^35^. In diabetic patients, we found several key mitochondrial changes in blood: cell-free mtDNA was increased in diabetes and in two separate studies of diabetic retinopathy and diabetic nephropathy, circulating mtDNA-CN was reduced in severe disease but inflammation and mtDNA damage were increased, with evidence of a disconnect between mtDNA content and mtDNA transcription^14,7, 36^. Whilst mtDNA-CN is a quantitative readout of mitochondrial genome abundance and can increase under conditions of chronic metabolic stress, the amount present may not always correspond with improved mitochondrial function and further studies are needed to understand the functional impact of higher mtDNA in the diabetic brain.

In contrast to our finding of higher mtDNA in the diabetic brain, in the periphery, much data suggests that lower circulating mtDNA-CN is associated with higher risk of several metabolic diseases including chronic kidney disease^18,37^ cardiovascular disease^15, 16^ and obesity^13^. However, there are very few studies looking at the long-term impact of metabolic disease on mtDNA-CN in target organs and tissues. Therefore, it remains unknown whether the pattern we are seeing in the brain is unique and differing from the periphery and the vascular system, or whether other organs affected in diabetes may show similar trends.

Our results raise the possibility that diabetic and non-diabetic AD are not biologically equivalent with respect to mitochondrial dysfunction and may involve distinct pathological processes. Importantly, mtDNA-CN represents only a single mitochondrial readout, and it is possible that other mitochondrial and metabolic markers may also differ according to diabetes status. This has implications for biomarker interpretation, since failure to account for diabetes may obscure biologically distinct processes within clinically defined AD cohorts. It may also have therapeutic implications, as interventions targeting metabolic or mitochondrial pathways may have different efficacy depending on the presence of comorbid diabetes. Future studies should therefore assess whether diabetic and non-diabetic AD differ in additional mitochondrial and metabolic markers, and whether these differences are associated with distinct responses to treatment. In this context, trials of metabolic therapies, including glucagon-like peptide-1 receptor agonists such as semaglutide, may be informative if outcomes are examined with reference to diabetes status. For instance, current trials of glucagon-like peptide-1 receptor agonists such as semaglutide could be examined for differential outcomes in AD patients with and without type 2 diabetes, given the opposing effects of AD and diabetes on mtDNA-CN that we describe here^38,39^. Public reporting in late 2025 indicated that EVOKE/EVOKE+ did not meet primary efficacy endpoints for slowing clinical progression overall and early subgroup analyses data suggest a similar lack of efficacy in patients with type 2 diabetes^40^. Even when overall efficacy is neutral, such large datasets provide an opportunity to test whether metabolic status, including diabetes, modifies biomarker trajectories or treatment response^40^.

A unique strength of the present study is the measurement of mtDNA-CN in the three brain regions from the same 64 donors, enabling assessment of whether mtDNA-CN levels are coordinated within individuals rather than evaluating each region in isolation. We have shown that diabetes is associated with a with a higher multiregional mtDNA-CN profile across the brain, whereas AD, especially in the absence of diabetes is associated with lower and more regionally variable mtDNA. These contrasting trends show that metabolic status influences AD and has potential therapeutic implications. As alterations to mitochondrial parameters appear system-wide rather than region-specific, therapeutic interventions may need to target multiple brain regions simultaneously for functional recovery. More broadly, the distinct multiregional pattern seen in D-AD versus ND-AD raises the possibility that metabolic status may influence mitochondrial therapeutic responsiveness, underscoring the need to consider comorbid diabetes when designing or evaluating mitochondria-targeted treatments for AD.

In summary, our findings show that AD is associated with cross-regional reduced mtDNA-CN, indicating a systemic loss of mitochondrial content, and supporting the broader view that impaired mitochondrial function contributes to AD pathology, potentially through reduced bioenergetic capacity. This effect was most evident in non-diabetic AD and was masked when diabetic AD cases were included. These findings support treating diabetes as an effect modifier rather than an optional covariate in AD studies. Wherever possible, diabetes-stratified analyses and formal diabetes-AD interaction testing alongside sensitivity analyses incorporating broader metabolic measures (e.g., obesity measures, glycaemic control, and medication exposure) could provide valuable information. Such an approach is likely to strengthen causal inference, improve biomarker interpretation, and prevent diabetes-related biology from being misattributed to AD, thereby improving both mechanistic understanding and therapeutic development.

## Abbreviations

AD: Alzheimer’s disease
mtDNA: mitochondrial DNA
mtDNA-CN: mitochondrial DNA copy number
NCI: non-cognitively impaired
ND-NCI: non-diabetic non-cognitively impaired
D-NCI: diabetic non-cognitively impaired
ND-AD: non-diabetic Alzheimer’s disease
D-AD: diabetic Alzheimer’s disease
PMI: post-mortem interval
PCA: principal component analysis
ICC: intraclass correlation coefficient
IQR: interquartile range
hMito: mitochondrial DNA target
hB2M: beta-2-microglobulin (nuclear DNA reference gene)

## ACKNOWLEDGEMENTS

We gratefully acknowledge funding for AK and the research undertaken for this work from Alzheimer’s Research UK (Interdisciplinary Research Grant ARUK-IRG2020A-001). Human brain tissue was obtained from the London Neurodegenerative Diseases Brain Bank (King’s College London) as part of the Brains for Dementia Research project.

## CONFLICT OF INTEREST

None

## SOURCES OF FUNDING

AK was supported by Alzheimer’s Research UK (Interdisciplinary Research Grant ARUK-IRG2020A-001). AS was supported by a China scholarship council award. ANM was partially supported by the PAS GRAS project, European Union grant agreement 101080329. RHS was supported by funding from P30AG072973.

## CONSENT STATEMENT

All donors had provided written informed consent for the use of postmortem brain for research purposes to the London Neurodegenerative Disease Brain Bank.

## SUPPLEMENTARY FILES

**Table S1:**
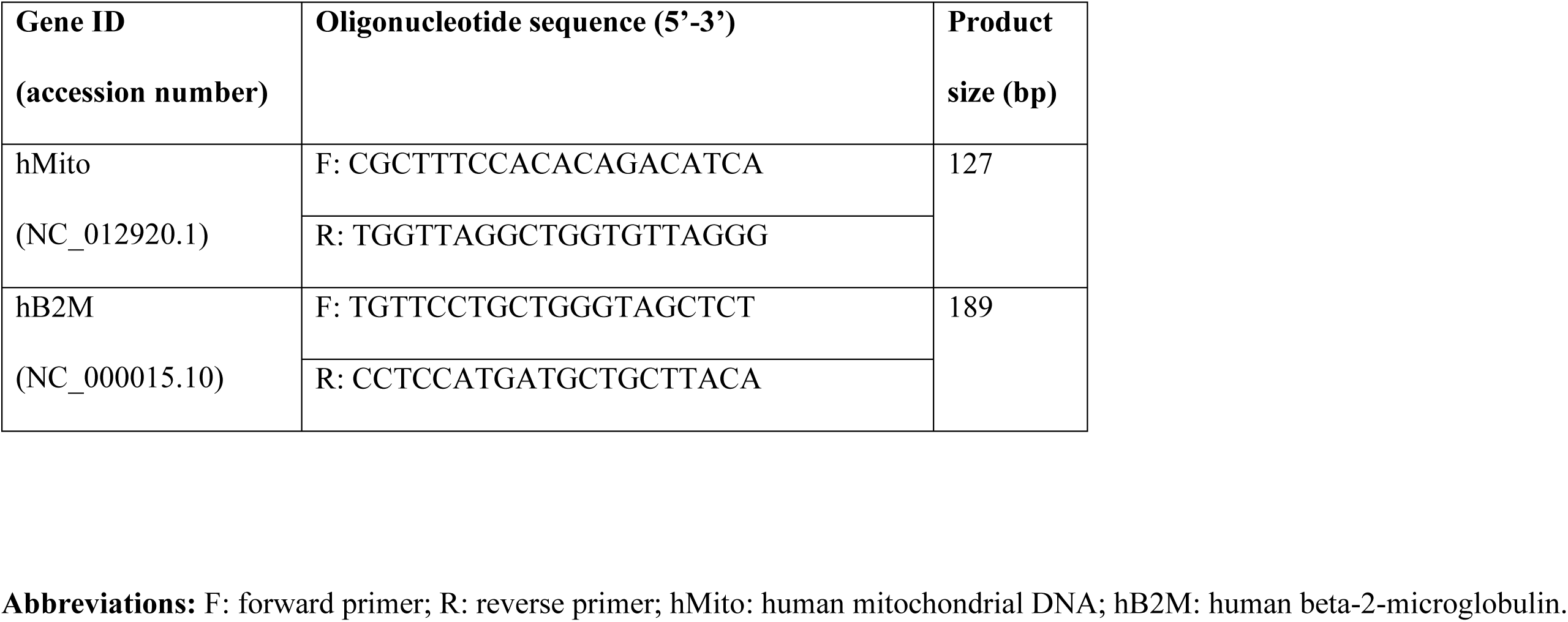
Oligonucleotide primers.

**Table S2.**
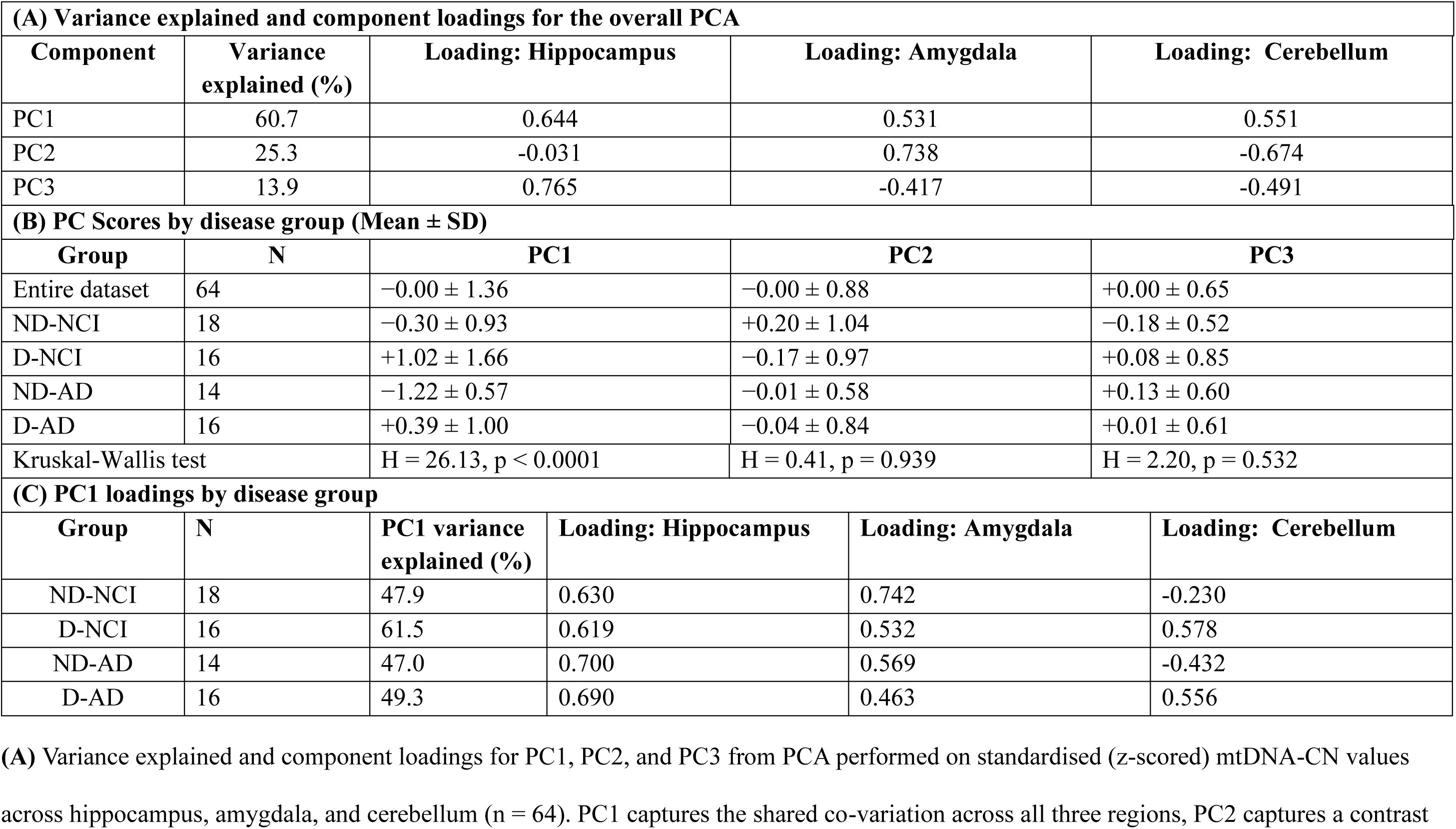
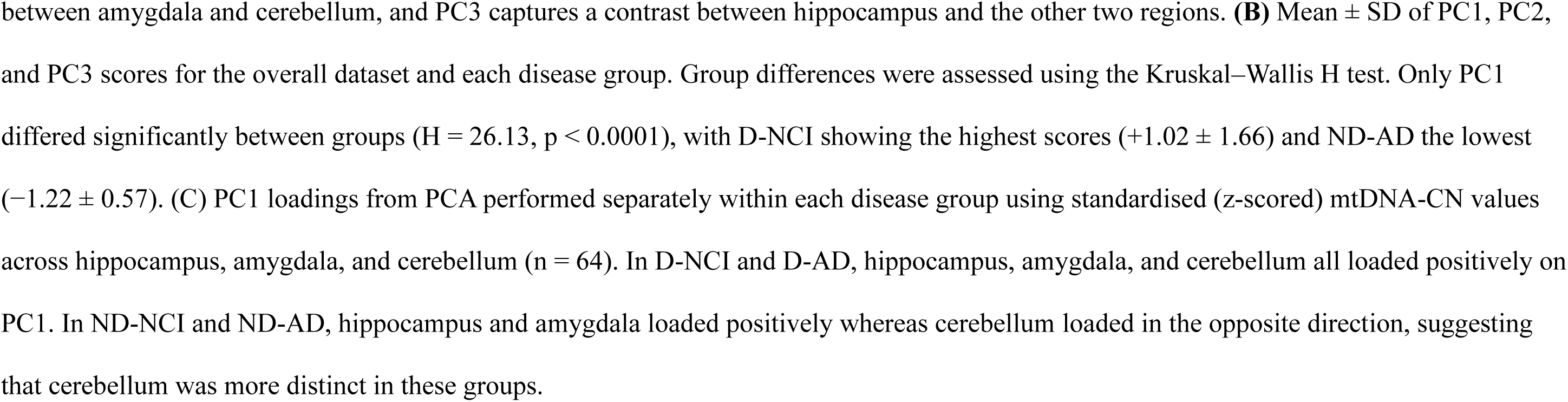
Principal component analysis of mtDNA-CN across three brain regions.

| (A) Variance explained and component loadings for the overall PCA |  |  |  |  |  |
| --- | --- | --- | --- | --- | --- |
| Component | Variance explained (%) | Loading: Hippocampus |  | Loading: Amygdala | Loading: Cerebellum |
| PC1 | 60.7 | 0.644 |  | 0.531 | 0.551 |
| PC2 | 25.3 | -0.031 |  | 0.738 | -0.674 |
| PC3 | 13.9 | 0.765 |  | -0.417 | -0.491 |
| (B) PC Scores by disease group (Mean ± SD) |  |  |  |  |  |
| Group | N | PC1 |  | PC2 | PC3 |
| Entire dataset | 64 | −0.00 ± 1.36 |  | −0.00 ± 0.88 | +0.00 ± 0.65 |
| ND-NCI | 18 | −0.30 ± 0.93 |  | +0.20 ± 1.04 | −0.18 ± 0.52 |
| D-NCI | 16 | +1.02 ± 1.66 |  | −0.17 ± 0.97 | +0.08 ± 0.85 |
| ND-AD | 14 | −1.22 ± 0.57 |  | −0.01 ± 0.58 | +0.13 ± 0.60 |
| D-AD | 16 | +0.39 ± 1.00 |  | −0.04 ± 0.84 | +0.01 ± 0.61 |
| Kruskal-Wallis test |  | H = 26.13, p < 0.0001 |  | H = 0.41, p = 0.939 | H = 2.20, p = 0.532 |
| (C) PC1 loadings by disease group |  |  |  |  |  |
| Group | N | PC1 variance explained (%) | Loading: Hippocampus | Loading: Amygdala | Loading: Cerebellum |
| ND-NCI | 18 | 47.9 | 0.630 | 0.742 | -0.230 |
| D-NCI | 16 | 61.5 | 0.619 | 0.532 | 0.578 |
| ND-AD | 14 | 47.0 | 0.700 | 0.569 | -0.432 |
| D-AD | 16 | 49.3 | 0.690 | 0.463 | 0.556 |

**(A)** Variance explained and component loadings for PC1, PC2, and PC3 from PCA performed on standardised (z-scored) mtDNA-CN values across hippocampus, amygdala, and cerebellum (n = 64). PC1 captures the shared co-variation across all three regions, PC2 captures a contrast
between amygdala and cerebellum, and PC3 captures a contrast between hippocampus and the other two regions. **(B)** Mean ± SD of PC1, PC2, and PC3 scores for the overall dataset and each disease group. Group differences were assessed using the Kruskal–Wallis H test. Only PC1 differed significantly between groups ( $H = 26.13$ , $p < 0.0001$ ), with D-NCI showing the highest scores ( $+1.02 \pm 1.66$ ) and ND-AD the lowest ( $-1.22 \pm 0.57$ ). **(C)** PC1 loadings from PCA performed separately within each disease group using standardised (z-scored) mtDNA-CN values across hippocampus, amygdala, and cerebellum ( $n = 64$ ). In D-NCI and D-AD, hippocampus, amygdala, and cerebellum all loaded positively on PC1. In ND-NCI and ND-AD, hippocampus and amygdala loaded positively whereas cerebellum loaded in the opposite direction, suggesting that cerebellum was more distinct in these groups.

**Table S3.**
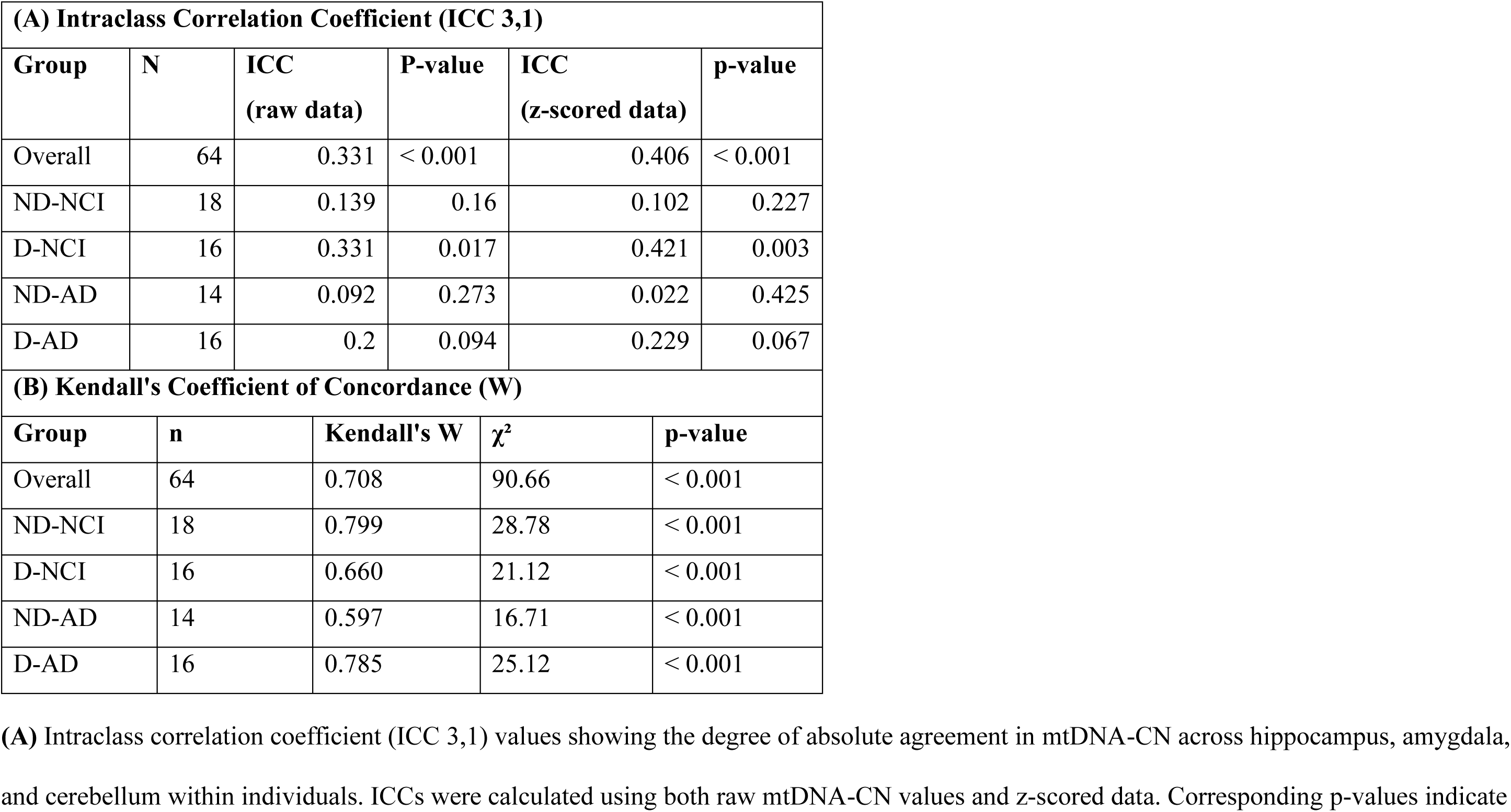
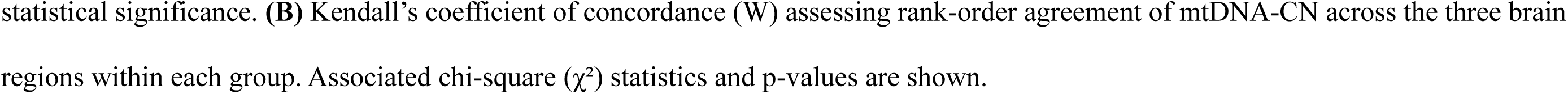
Intraclass correlation coefficient (ICC) and Kendall’s coefficient of concordance (W) for mtDNA-CN in three brain regions.

| **(A) Intraclass Correlation Coefficient (ICC 3,1)** |  |  |  |  |  |
| --- | --- | --- | --- | --- | --- |
| **Group** | **N** | **ICC**<br>**(raw data)** | **P-value** | **ICC**<br>**(z-scored data)** | **p-value** |
| Overall | 64 | 0.331 | < 0.001 | 0.406 | < 0.001 |
| ND-NCI | 18 | 0.139 | 0.16 | 0.102 | 0.227 |
| D-NCI | 16 | 0.331 | 0.017 | 0.421 | 0.003 |
| ND-AD | 14 | 0.092 | 0.273 | 0.022 | 0.425 |
| D-AD | 16 | 0.2 | 0.094 | 0.229 | 0.067 |
| **(B) Kendall's Coefficient of Concordance (W)** |  |  |  |  |  |
| **Group** | **n** | **Kendall's W** | **$\chi^2$** | **p-value** |  |
| Overall | 64 | 0.708 | 90.66 | < 0.001 |  |
| ND-NCI | 18 | 0.799 | 28.78 | < 0.001 |  |
| D-NCI | 16 | 0.660 | 21.12 | < 0.001 |  |
| ND-AD | 14 | 0.597 | 16.71 | < 0.001 |  |
| D-AD | 16 | 0.785 | 25.12 | < 0.001 |  |

**(A)** Intraclass correlation coefficient (ICC 3,1) values showing the degree of absolute agreement in mtDNA-CN across hippocampus, amygdala, and cerebellum within individuals. ICCs were calculated using both raw mtDNA-CN values and z-scored data. Corresponding p-values indicate
statistical significance. **(B)** Kendall's coefficient of concordance (W) assessing rank-order agreement of mtDNA-CN across the three brain regions within each group. Associated chi-square ( $\chi^2$ ) statistics and p-values are shown.

**Table S4.**
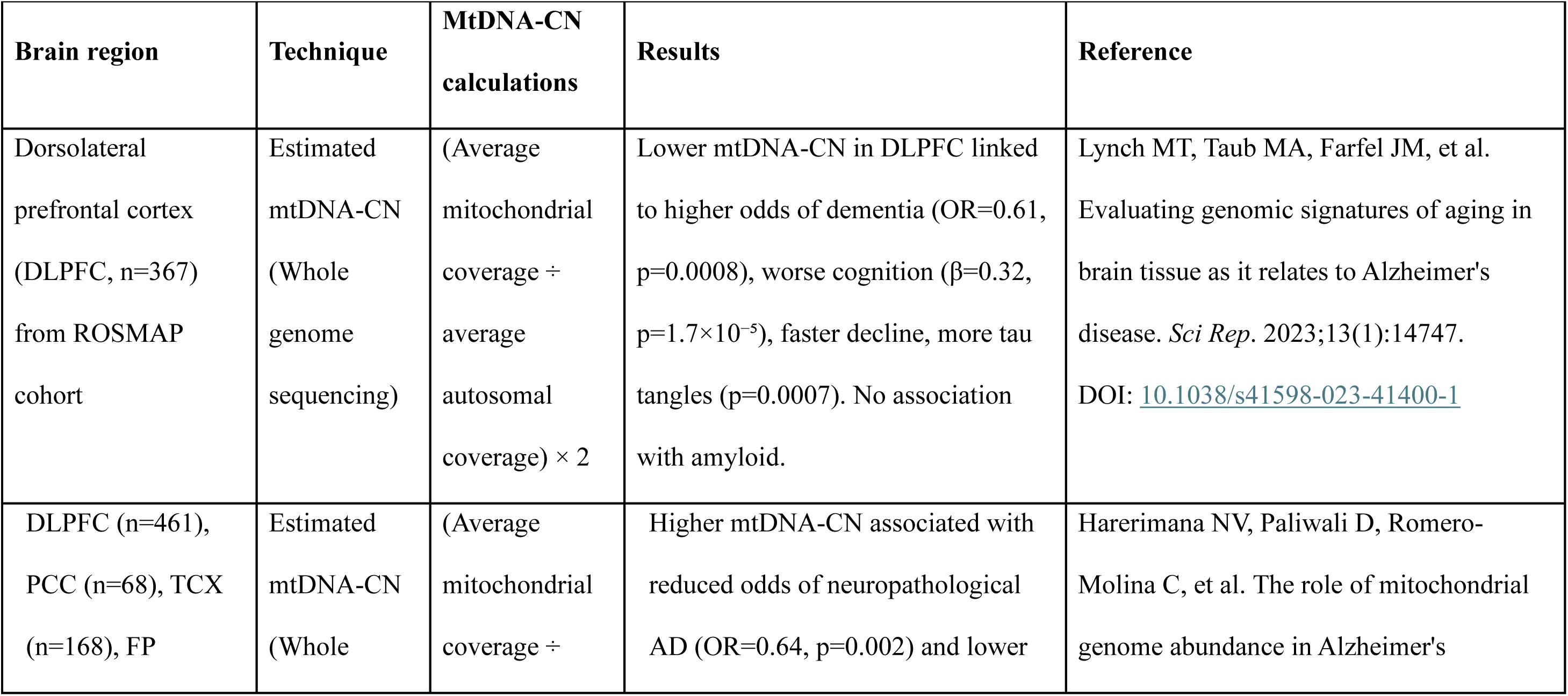
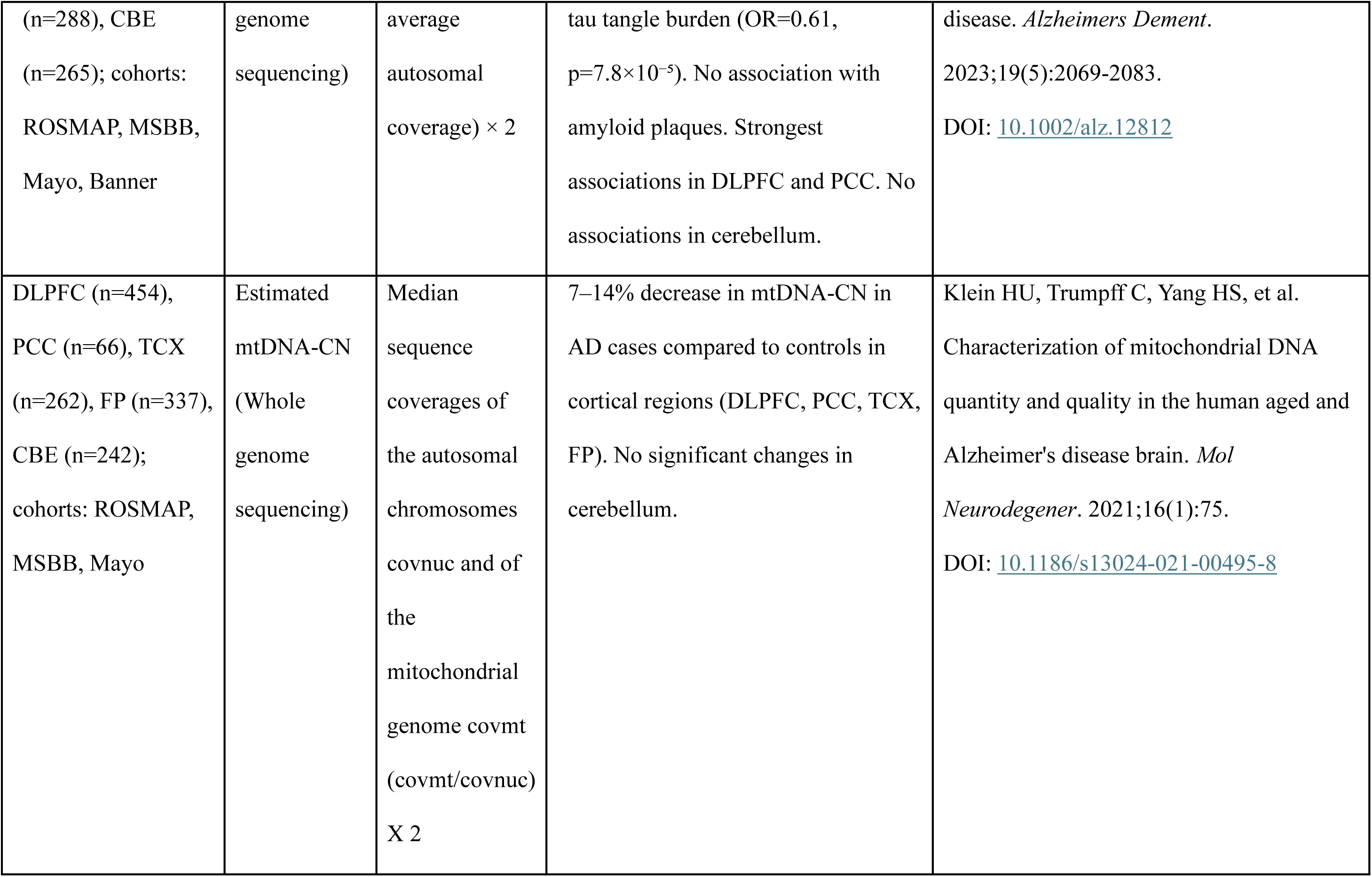
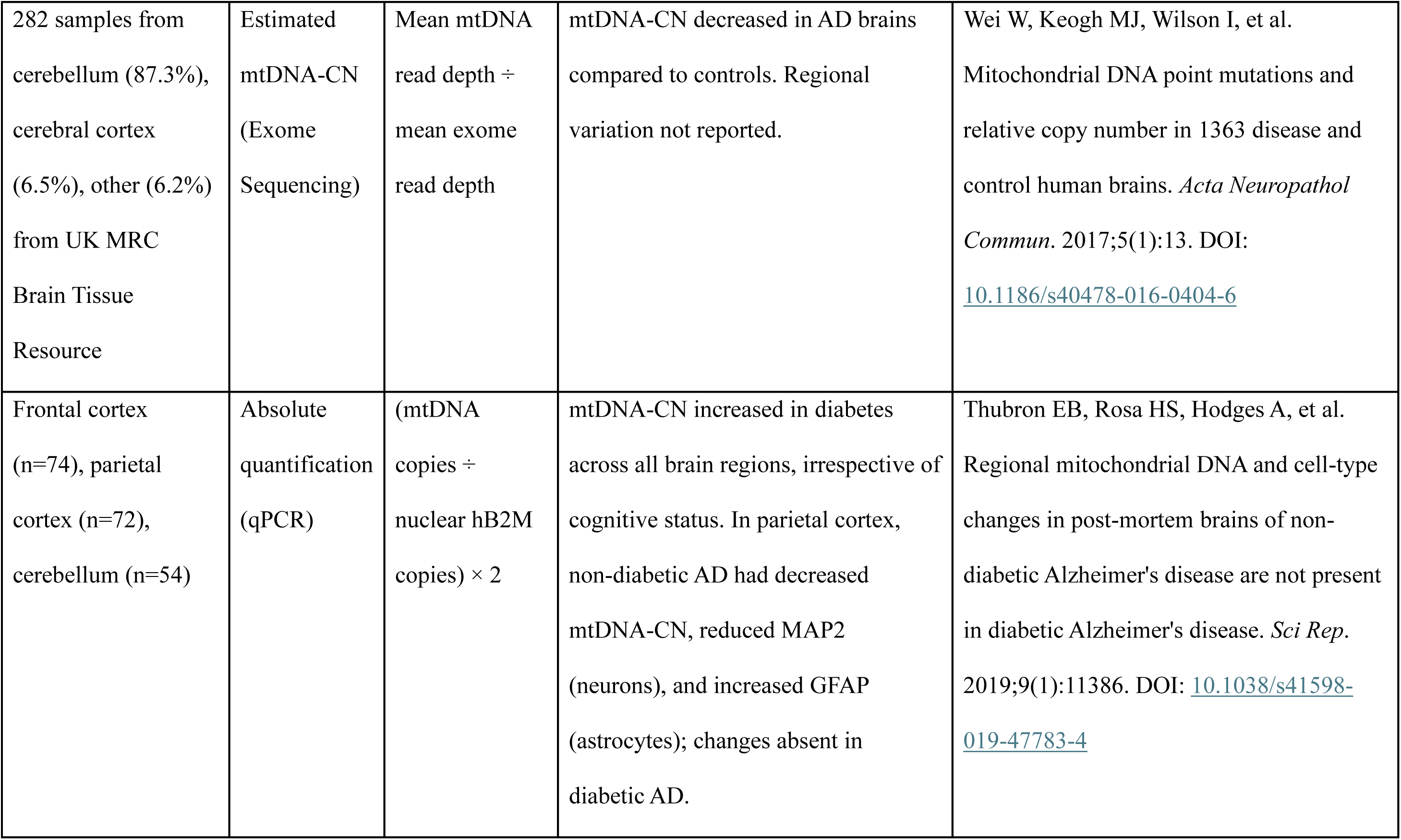
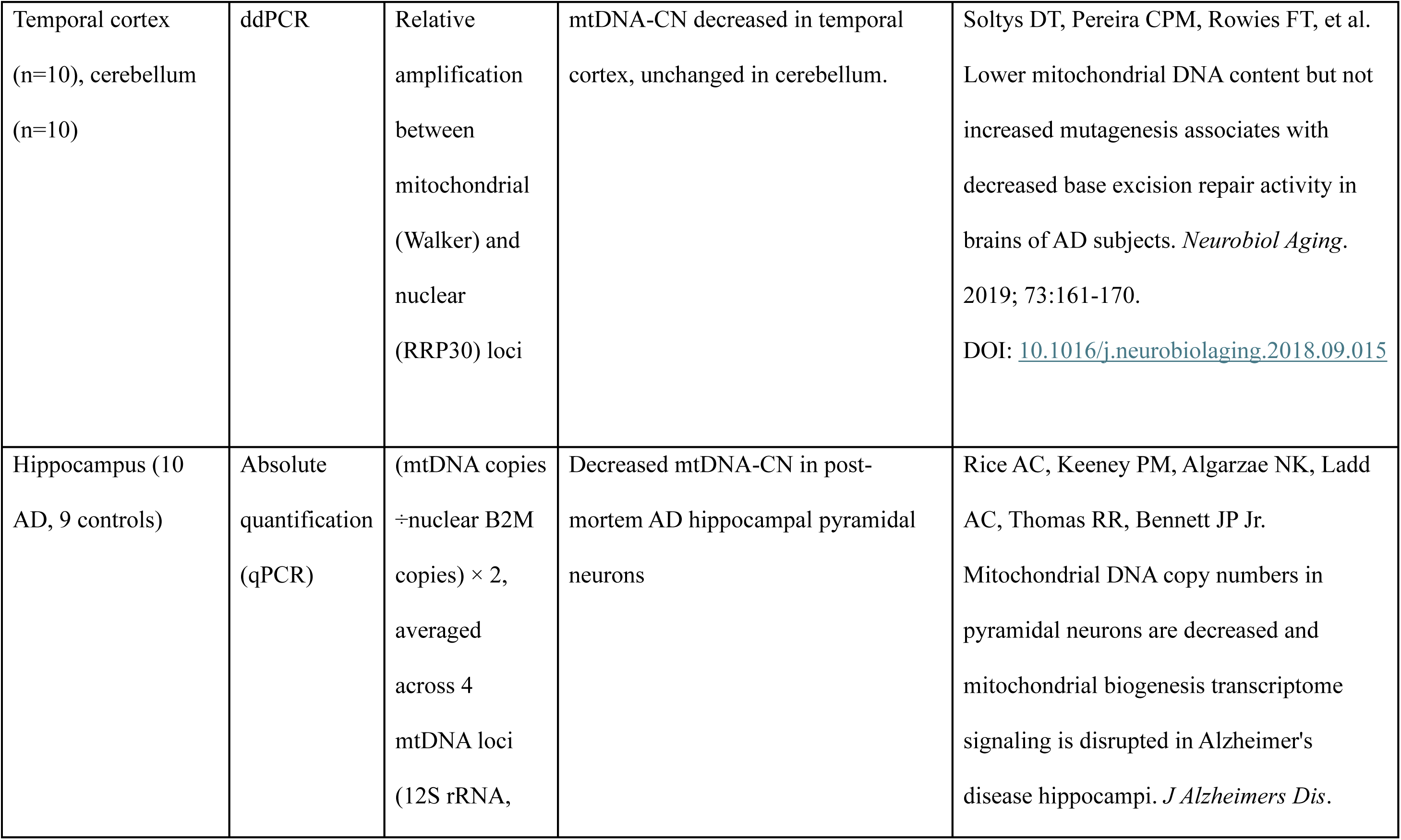
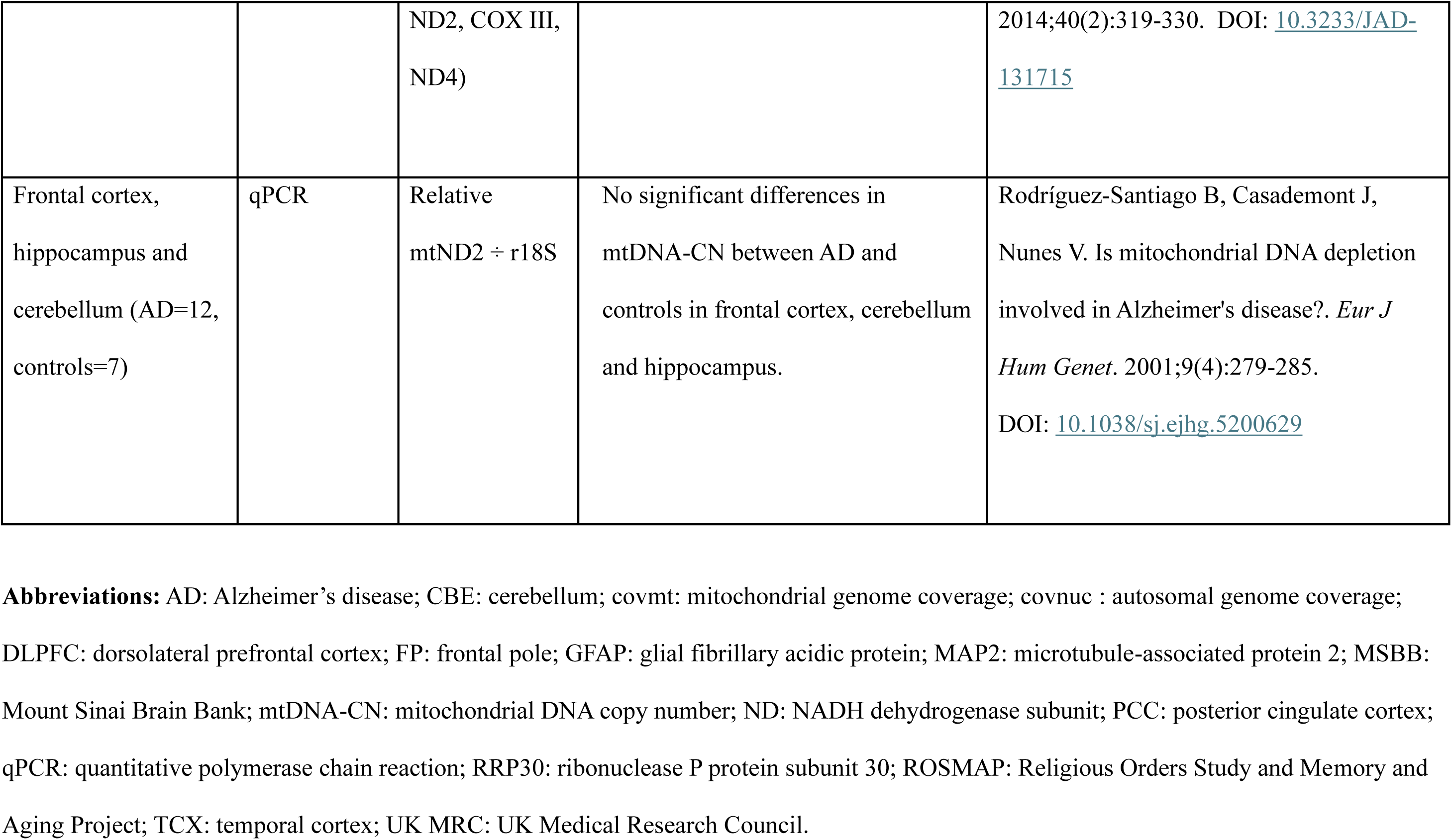
Summary of published studies evaluating mitochondrial DNA copy number in Alzheimer’s disease using human post-mortem brain tissue.

## Notes

### Competing Interest Statement

The authors have declared no competing interest.

